# Addressing the problem of lysine glycation prediction in proteins via Recurrent Neural Networks

**DOI:** 10.1101/2024.08.12.607666

**Authors:** Ulices Que-Salinas, Dulce Martinez-Peon, Gerardo Maximiliano Mendez, P. Argüelles-Lucho, Angel D. Reyes-Figueroa, Christian Quintus Scheckhuber

**Affiliations:** Centro de Ciencias de la Tierra, Universidad Veracruzana, Xalapa 91090, VER, México; Tecnológico Nacional de México/ Instituto Tecnológico de Nuevo León, Departamento de Ingeniería Eléctrica y Electrónica, Av. Eloy Cavazos 2001, Guadalupe, NL, México; Tecnológico Nacional de México, Instituto Tecnológico de Veracruz, Calzada Miguel Ángel de Quevedo 2779, Veracruz, Ver 91860, México; Consejo Nacional de Ciencia y Tecnología, Av. Insurgentes Sur 1582, Col. Crédito Constructor, Benito Juárez, México City 03940, DF, México; Centro de Investigación en Matemáticas Unidad Monterrey, Parque de Investigación e Innovación Tecnológica (PIIT), Av. Alianza Centro No. 502, Apodaca 66628, NL, México; Tecnológico de Monterrey, Escuela de Ingeniería y Ciencias, Av. Eugenio Garza Sada 2501 Sur, Col. Tecnológico, Monterrey 64700, NL, México

**Keywords:** amino acids, classification, glycation, lysine, protein sequence, recurrent neural networks

## Abstract

A distinguishing feature of the metabolic disorder diabetes involves elevated damage to cellular proteins. The primary form of alteration arises from the chemical interaction between glycating agents such as methylglyoxal and proteinaceous arginine/lysine residues, causing structural and functional disruptions in target proteins. In this study, a curated version of the CPLM database to implement a recurrent neural network strategy for the classification of lysine glycation has been utilized. By using one physical property for the characterization of amino acids next to lysine sites (i.e., isoelectric point), it was possible to obtain a 59.6% accuracy for correctly predicting lysine glycation. When two properties were combined, i.e., mass and torsion angle, the accuracy increased to 59.9%. Overall, this approach can aid the task of narrowing down possible sites of lysine glycation in protein targets for further analysis.

## 1. Introduction

Proteins are constructed from a rather small set of unique amino acids. The roster of amino acids comprises 20 ’standard’ ones, along with a few more esoteric proteinogenic variants, such as selenocysteine and pyrrolysine (Ho et al., 2021). This allows for an astronomical range of individual potential sequences, even in relatively short proteins. In addition to this staggering diversity, the plethora of post-translational modifications that amino acids can undergo should be considered. These modifications introduce layers of regulation and control, encompassing an array of processes leading to adducts. Notable examples include acetylation, phosphorylation, methylation, and ubiquitination, among several others (Müller, 2018). The post-translational modification of specific amino acids can occur enzymatically or through non-enzymatic means. For instance, glycosylation, vital for protein sorting, secretion and cellular recognition, is accomplished by N- and O-glycosyltransferases and other enzymes (Williams & Thorson, 2008). Another critical example involves the reversible modification of histones by histone acetylases and deacetylases, a fundamental process for the coordinated regulation of gene expression (Y. Zhang et al., 2021).

In contrast, glycation is a predominantly non-enzymatic process, entailing the reaction of sugars (e.g., glucose, fructose) and sugar-derived compounds with biologically significant molecules, such as nucleic acids, lipids, and proteins (Rabbani & Thornalley, 2015). First, a Schiff base between the target and the glycating agent is formed which is rearranged to yield an Amadori product. Typically, these reactions culminate in the formation of advanced glycation end-products (AGEs), most of which exert detrimental effects and irreversibly compromise the function of the target molecules (Ahmed et al., 2005; Oya et al., 1999).

Within proteins, the side chains of lysine and arginine emerge as primary targets for AGE formation (Mercado-Uribe et al., 2020; Rabbani & Thornalley, 2021). Methylglyoxal (MGO), a highly reactive glycating compound, ranks among the most prolific culprits. It arises as a toxic by-product during metabolic processes, such as glycolysis, in which it is produced by spontaneous elimination of phosphate from the glycolytic intermediate dihydroxyacetone phosphate (Phillips & Thornalley, 1993). Under physiologically normal circumstances, cellular MGO levels remain relatively low, typically hovering around 0.3–6 µM (Rabbani & Thornalley, 2014). This is upheld by dedicated enzymatic defense systems, such as glyoxalase I and II, aldose reductases, and low-molecular-weight scavengers. However, in certain pathological conditions (e.g., diabetes, neurodegeneration, cancer), and in aging cells and tissues, MGO can pose challenges to cellular viability due to heightened production or hindered removal (Kumar Pasupulati et al., 2016; Schalkwijk & Stehouwer, 2020). It is important to note that MGO-mediated protein modifications can play a pivotal role in various signaling processes and gene regulation. This has been substantiated through dedicated studies, often conducted in simple eukaryotic model systems known for their experimental tractability (Scheckhuber, 2019).

Despite extensive research on the importance of MGO binding to specific amino acids in target proteins, it has become evident that there is no straightforward consensus sequence for reliably predicting potential glycation sites (Sjoblom et al., 2018). Still, several promising approaches have been made that allow for the prediction of potential lysine glycation sites. Examples include GlyNN (Johansen et al., 2006), which employs an artificial neural network (ANN) (Rabuñal & Dorado, 2006) to predict lysine glycation, and BPB_GlySite (Ju et al., 2017), PreGly (Y. Liu et al., 2015), PredGly (Yu et al., 2019), Gly–PseAAC (Y. Xu et al., 2017), Glypre (Zhao et al., 2017), iProtGly–SS (Islam et al., 2018), GlyStruct (Reddy et al., 2019) as well as BERT-Kgly (Y. Liu et al., 2022) and iGly-IDN (Jia et al., 2024). These methods employ approaches such as bi-profile Bayes features extraction, position-specific amino acid propensity, transformers (BERT)-based models and models trained with support vector machine (SVM) classifiers. Traditionally, lysine modification has been the focus of extensive research, leading to the availability of a comprehensive database of lysine modifications, such as PLMD (H. Xu et al., 2017), based on CPLM (Z. Liu et al., 2014) and CPLA 1.0 (Z. Liu et al., 2011). For the detection of arginine-dependent glycation a predictor based on artificial neural networks that utilizes short peptides that were subjected to controlled in vitro glycation, yielding a small but high-quality data set was recently published (Que-Salinas et al., 2022; Sjoblom et al., 2018).

It is relevant to narrow down potential sites of protein modification, as the experimental demonstration of specific amino acid modifications can be prohibitively expensive in terms of time, resources, and labor, particularly in larger proteins housing numerous potential glycation sites.

The approach presented here uses a high-quality protein lysine modification data set to employ a recurrent neural network (RNN) to classify lysine glycation. It shows that using one physical property for peptide characterization, isoelectric point, a 59.6% accuracy for lysine glycation is obtained. Additionally, when combining the two properties, mass and torsion angle, the accuracy increases to 59.9%.

## 2. Materials and methods

### 2.1 General outline of the experimental approach

Figure 1 shows the methodology that was followed. First, redundant information from the CPLM 4.0 database (Z. Liu et al., 2014; W. Zhang et al., 2022) was removed by utilizing CD-HIT (Huang et al., 2010) with a 30% cut-off. From this, 6830 peptides, each a sequence of 31 amino acids, were obtained.

**Figure 1.**
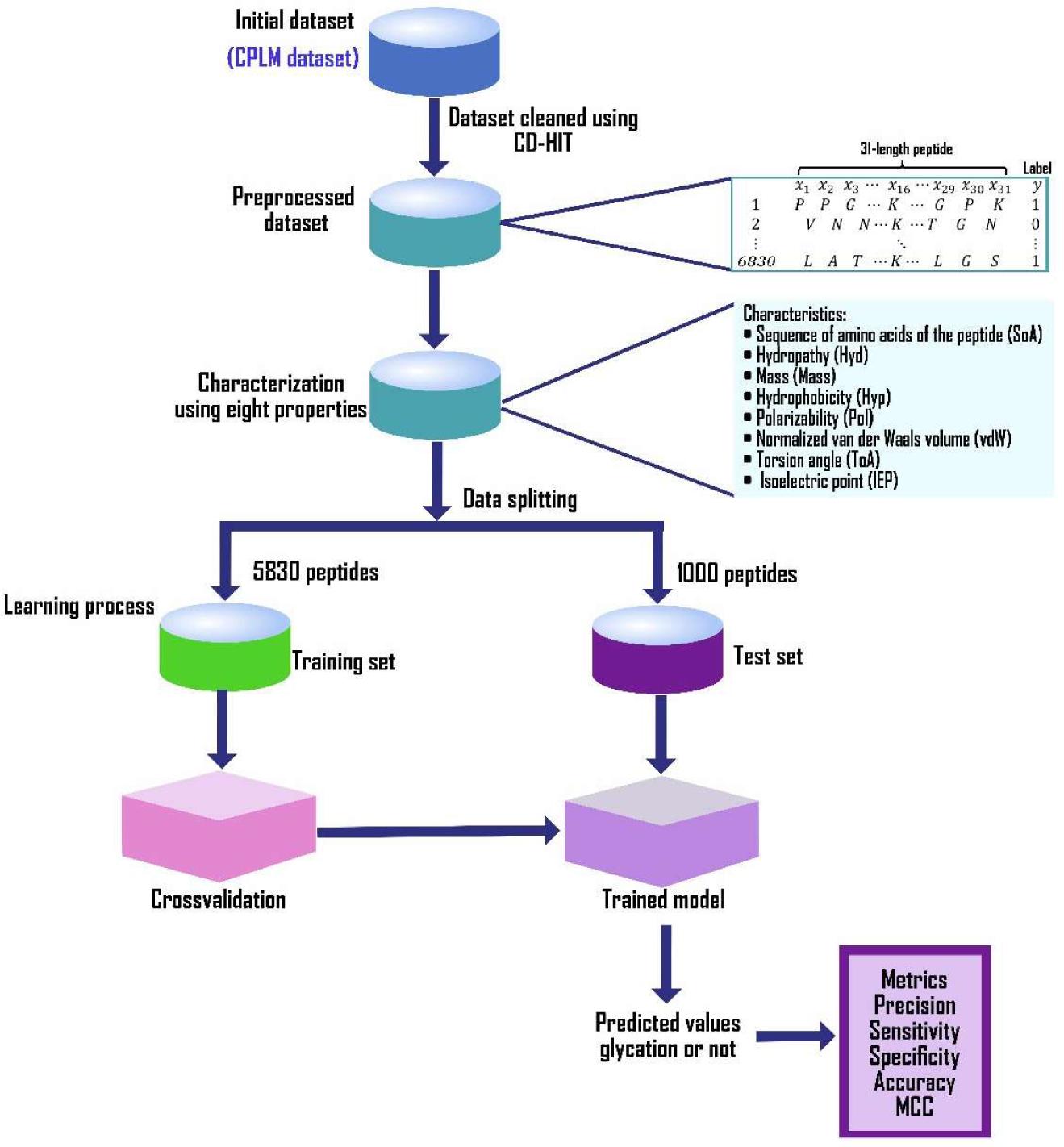
Flowchart of the steps to classify glycation of lysine. The data collected from the CPLM 4.0 database were processed using the CD-HIT tool, forming arrays of vectors characterized by the physical properties of the amino acids. The processed database was divided into three sets (training, validation, and testing) to perform the RNN learning process and test the results obtained through a series of metrics.

The sequences of amino acids are characterized by up to eight different physical properties: the proper sequence of amino acids, i.e., the structure of the amino acid sequence (SoA) of the peptide, hydropathy (Hyd), mass (Mass), hydrophobicity (Hyp), polarizability (Pol), normalized van der Waals volume (vdW), torsion angle (ToA), and isoelectric point (IEP) (Table S1 (Que-Salinas et al., 2022). These amino acid sequences were used to feed a numerical algorithm based on Recurrent Neural Networks (RNN), which was utilized as a model for the intelligent recognition process of the presence of glycation in lysine. As the final stage of the methodology, the trained model was validated on an independent test set. Therefore, the following metrics were determined as quantitative parameters to evaluate the performance of the model: (i) accuracy, the ratio between the correct predictions for both true glycated and true non-glycated proteins; (ii) precision, a ratio based on the prediction of true glycated proteins and the sum of true and false positive glycation prediction, (iii) sensitivity, a ratio for the prediction of true glycated proteins based on true positive and false negative glycation prediction, (iv) specificity, a ratio between the prediction of true non glycated proteins based on the true and false prediction of non-glycation, and (v) the Matthews Correlation Coefficient (MCC), a metric that measures the correlation between the model and the real system (Jurman et al., 2012; Y. Liu et al., 2022). Although these metrics have values ranging from 0 to 1, representing a 1 that the model correctly predicts 100% of the real values, it should be considered that in the case of MCC, the values range from -1 to 1, which means perfect misclassification or classification, respectively. At the same time, MCC=0 is the expected value for the “coin tossing” classifier. It is worth mentioning that MCC is the only metric that considers whether the classifier was able to correctly predict both the majority of positive and negative data instances (Jurman et al., 2012).

### 2.2 Characterization scheme

First, the symbolic letters that represent the amino acids of the 6830 peptides were replaced by a numerical value linked to a particular physical property (see Tab. S1). For each peptide, eight different vectors of 31 numerical values were generated, each one of them representing one of the physical properties previously discussed. Thus, the vectors maintain the same order of the amino acids but are now represented by the numerical value corresponding to one of these properties. Therefore, each of the 6830 peptides used for this study will be represented by eight vectors of 31 numerical values.

For our numerical model learning process, the samples were divided in the following way: 4830 sequences for training, 1000 peptides were used for the internal learning validation process, and the remaining 1000 were used to verify the performance of the model through an independent test. For the training step, a recurrent neural network (RNN) using the characterized data with the eight properties SoA, Hyd, Mass, Hyp, Pol, vdW, ToA, and IEP was obtained (Que-Salinas et al., 2022).

### 2.3 Recurrent Neural Network (RNN) architecture

The structure of the RNN used was of the long short-term memory (LSTM) type. It contains an input layer, a hidden layer, and an output layer, as shown in Figure 2. The output layer has two neurons, one to indicate the probability of the presence of glycation on the central amino acid (i.e., lysine) and the other for the opposite. The outcome of the RNN was determined by considering which of the two options presents a higher probability, repeating the process a total of 20 times to ensure the reproducibility of the model. The hidden layer is made up of six distinct layers. The first contains 64 neurons; each neuron is the unity of LSTM with a tangent hyperbolic function and a linear regularized l2 type, followed by a dropout layer. After that, there is another layer of 32 LSTM units with the same configuration as those used in the first hidden layer. This is followed again by a second dropout layer that contributes to forgetting the adjusted weights to avoid overtraining. After that, a dense layer is highly connected, composed of 16 neurons with a rectified linear unit (ReLU) activation function followed by a dropout layer. Finally, the input layer consists of an arrangement of neurons that receives the internal values of an array of 31X2 elements for cases 1 and 2 and 31X8 for case 3 (see below for the description of the cases).

**Figure 2.**
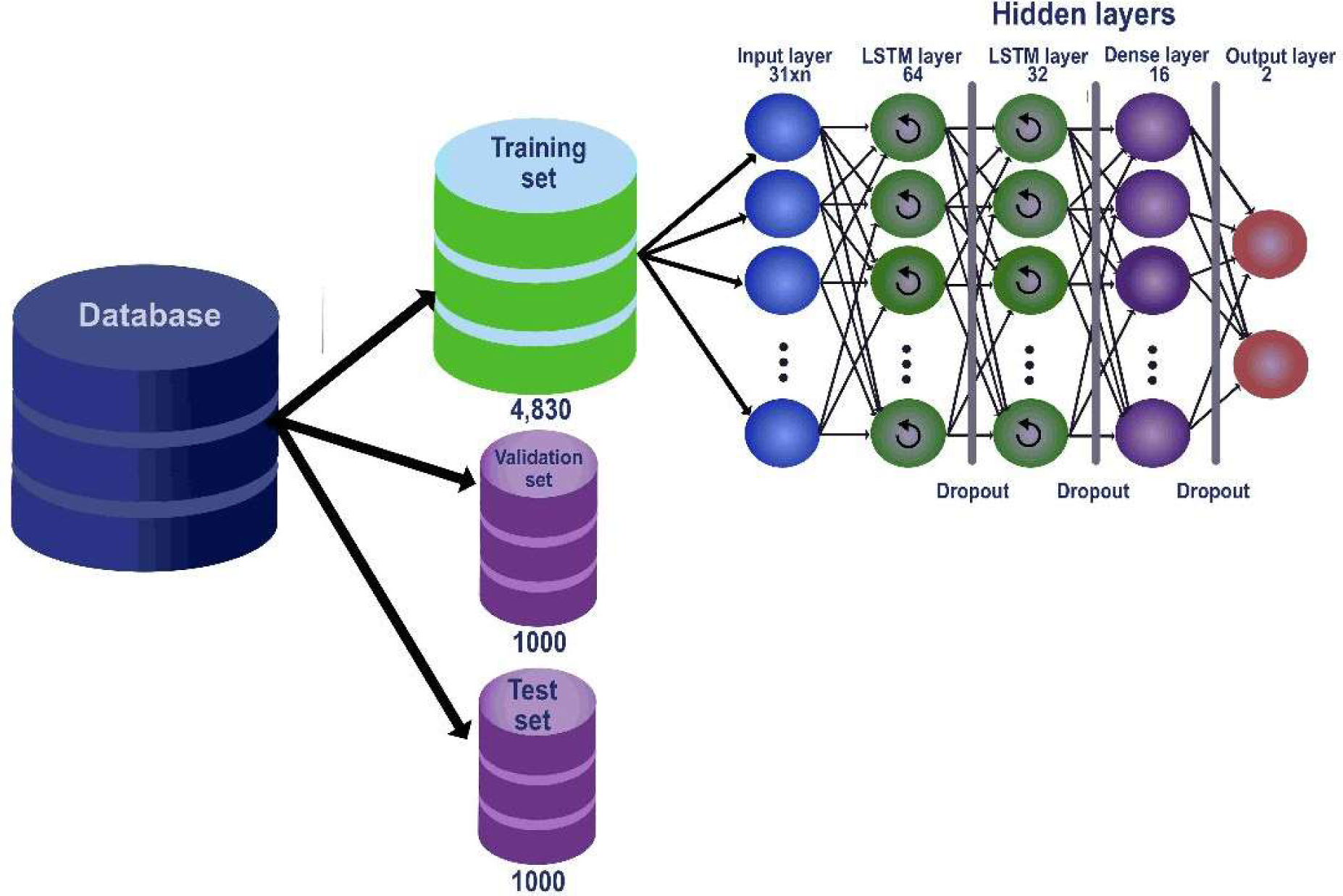
Structure of the RNN. Once the database has been divided into its three sets, the training set is used for the learning process of the RNN. This consisted of two hidden LSTM type recurrence layers and a dense layer followed by the output layer of the results. This process is shown from left to right, indicating at the top of each RNN layer the number of neurons that compose it.

For the RNN a sparse categorical cross entropy cost function with an ADAM type optimizer was used, and the accuracy metric to monitor the internal results was determined (Chandriah & Naraganahalli, 2021). For the training, a batch size of 64 was applied, with several training epochs that depended on an early stopping algorithm: if the accuracy after 20 epochs does not increase compared with the validation set, the training stops, and the epoch that contains the better results is maintained. Due to the early stopping algorithm, the number of training epochs varied approximately between 30 and 120.

Subsequently, the learning of the RNN was achieved with the training dataset using a set of validation to monitor the performance and to follow the end of the process, reviewing for this purpose the values of some metrics (such as accuracy) throughout the training evolution. Three different cases were defined to test the efficiency of the model and achieve the objective of this study:

Case 1: For each of the eight properties, a matrix of length 31X2 was built, where 31 is the length of the peptide and contains the obtained values of each property for each amino acid of the peptide, the mass, for example. This was duplicated for the second column. The approach allows for the analysis of the effect of each of the physical properties of the amino acids separately.

Case 2: Combinations of two properties for the input of the RNN. The input size was 31X2, where the first column belongs to the values of one property and the second column belongs to the values of a second property; for the same peptide, the total number of combinations was 28. This second case allows for determining the weight of the combinations of physical properties in the glycation process.

Case 3: Total number of properties for the input of the RNN. The input size matrix was 31X8, where each column was assigned to one property.

## 3. Results

Our RNN strategy was applied to the processed database that is outlined in section 2.1. The goal was to predict the presence or absence of lysine glycation in the peptides of the independent test set. The results of this analysis consist of the quantitative values corresponding to the five metrics described in section 2.1.

Consider case 1 for the property IEP: the test set is composed of a list of 1000 peptides, where for each one of these, the RNN will assign a probability that there is glycation and another one that there is no glycation, totaling between them a 100% probability. Thus, a list of labels corresponding to the answer of whether there is glycation (represented by a 1) or no glycation (represented by a 0) has been built, where, on the respective resulting quantitative values for the entire test set, each of the five metrics defined in section 2.1 were calculated. If the same process is run a total of 20 times and the values obtained for each metric are averaged, a more reliable value of the RNN performance can be obtained, which will be linked to the probability of glycation or non-glycation on the respective peptide. This same procedure was applied for the rest of the properties, as well as for all the sub-cases of case 2 and case 3. In this way, a global evaluation and comparison of the multiple subcases studied in this work can be obtained. The label with the highest probability of glycation will be considered further.

Figs. 3, 4, S1-S4, and table S2 show the results obtained for case 1 with respect to each of the metrics studied. It can be observed that the IEP is the physical property that yields the highest values for the metrics Acc (0.596), Pre (0.584) and Spe (0.553). For the case of sensitivity, the highest value was 0.835, corresponding to the torsion angle; similarly, for the MCC, the highest value reached was 0.196 corresponding to the mass. Regarding the lowest values, for accuracy and MCC, the respective values of 0.576 (torsion angle / SoA) and 0.172 (hydrophobicity / SoA) were obtained. A detailed analysis of the precision, sensitivity, specificity metrics and the MCC for case 1 is shown in Figs. S1 to S4.

**Figure 3.**
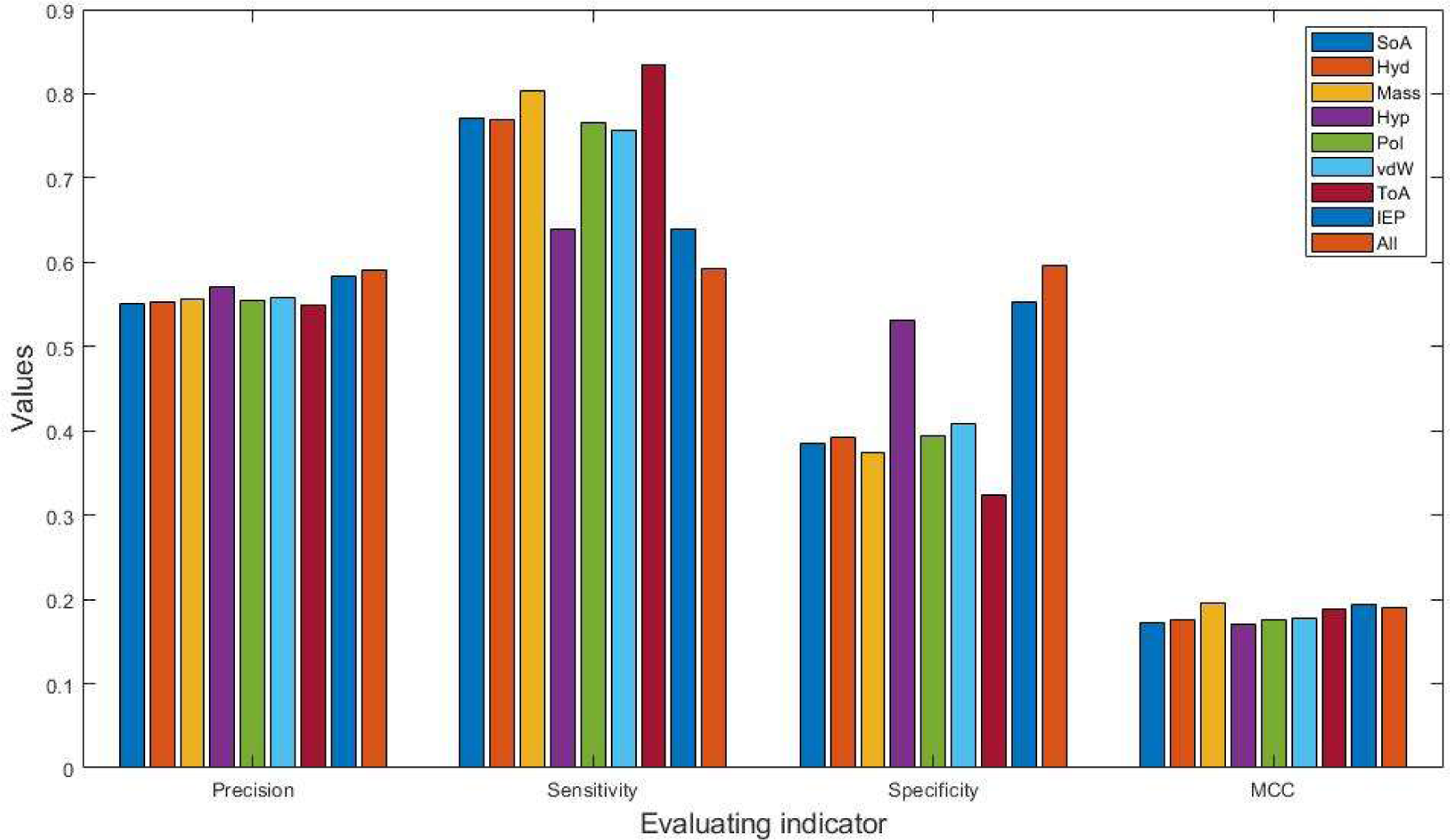
Performance metrics for case 1 and case 3. The four histograms correspond to the application of the metrics (Evaluating indicator): precision, sensitivity, specificity and MCC. Each histogram shows the results for case 1 (where all eight properties were tested individually) in its first eight columns and for case 3 in the last column (where all eight properties were tested together). Mean values of each metric are shown.

**Figure 4.**
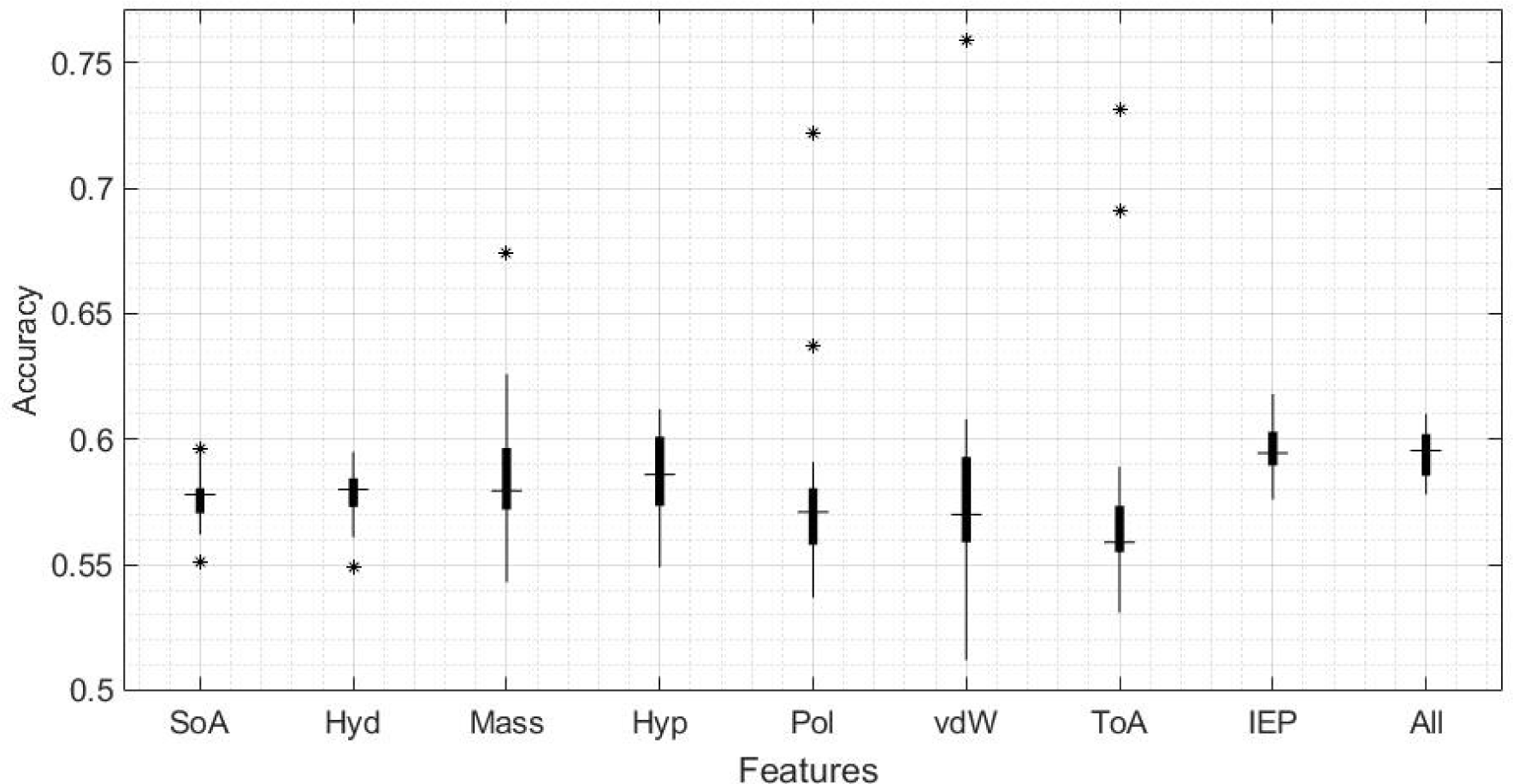
Detailed analysis of the accuracy metric for case 1 and case 3. The x-axis lists the eight properties analyzed in case 1 plus the combination of the eight properties for a single analysis corresponding to case 3 (all properties combined). The y-axis shows the values achieved for the accuracy metric. Median values are indicated by horizontal lines, the * symbol denotes outliers.

For case 2, the combination of both mass and torsion angle presented the highest accuracy value with 0.599 and in precision with a value of 0.583, respectively (Fig. 5, 6, and Tab. S3). It is noteworthy to point out that Mass was the property that presented the highest values in the metrics applied after IEP for case 1 and that it was found in two of the five best combinations of case 2. Other properties that showed a notable effect on the performance of the RNN for this case were IEP, SoA and Hyp. A detailed analysis of the accuracy for all property combinations of case 2 is shown in Fig. S5. Overall, for case 2, only a slight improvement is achieved when compared to case 1. Thus, we cannot state that combining three or more physical properties will increase the performance of the RNN substantially, considering the contrast between the increase in computational complexity versus the improvement in performance.

**Figure 5.**
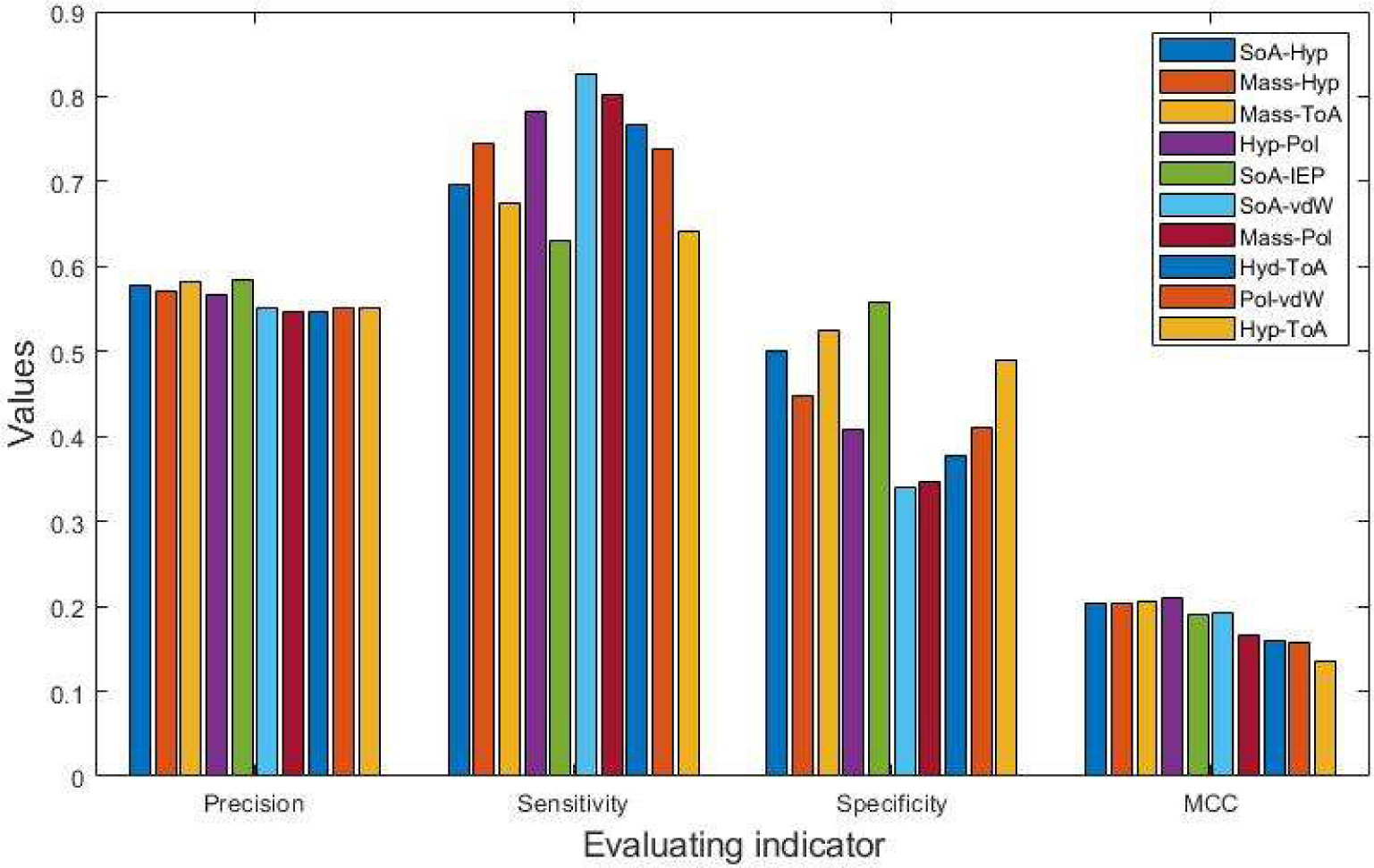
Performance metrics for case 2. The four histograms correspond to the application of the metrics (Evaluating indicator): precision, sensitivity, specificity and MCC. Each histogram shows the results for the strongest (SoA+Hyp, Mass+Hyp, Mass+ToA, Hyp+Pol, SoA+IEP) and weakest (SoA+vdW, Mass+Pol, Hyd+ToA, Pol+vdW, Hyp+ToA) candidates of the accuracy metric. Mean values of each metric are shown.

**Figure 6.**
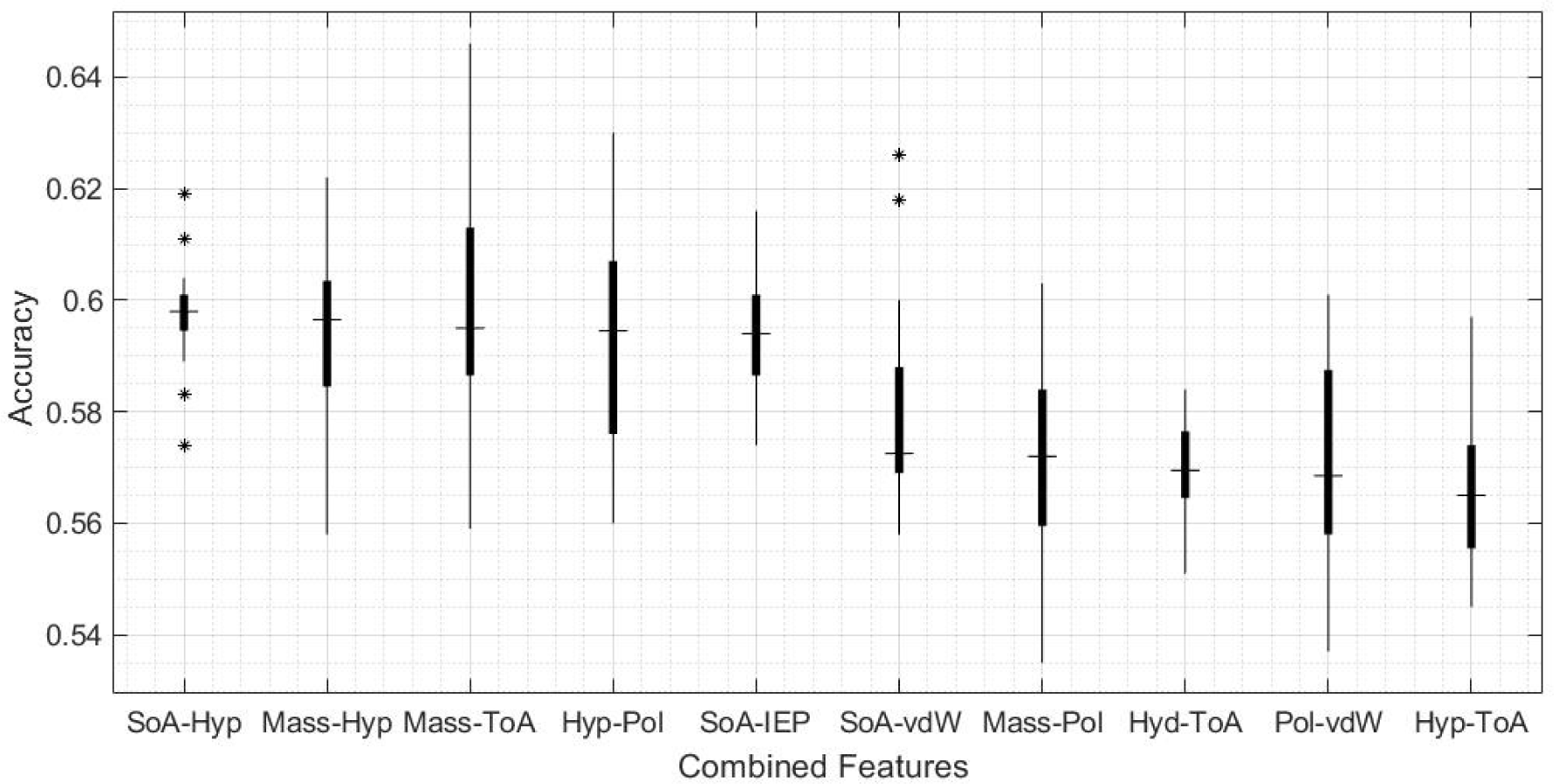
Detailed analysis of the accuracy metric for selected case 2 candidates. Box plots for the strongest (SoA+Hyp, Mass+Hyp, Mass+ToA, Hyp+Pol, SoA+IEP) and weakest (SoA+vdW, Mass+Pol, Hyd+ToA, Pol+vdW, Hyp+ToA) candidates. Median values are indicated, the * symbol denotes outliers.

For case 3 (Fig. 3, 4, Tab. S4), it is observed that the values obtained for all metrics were like those reported for both case 2 and case 1. As already mentioned, this infers that adding more physical properties does not necessarily improve accuracy but can result in the opposite, generating more complexity to the learning of the RNN, so this requires greater complexity in the RNN and, therefore, greater computational power. This is the reason why, in the present work, combinations of three or more physical properties are not included, considering that such cases imply the execution of many studies sub-cases (corresponding to each of the possible combinations), which, in preliminary tests, did not show an acceptable improvement in the performance of the RNN.

## 4. Discussion

In this work, a tool for predicting the presence of glycated lysine residues in protein sequences is presented. This was approached by identifying which of the physical properties of amino acids adjacent to the site of modification is the most relevant for the glycation process. The developed algorithm is based on an RNN and shows that of all these properties, the one that shows the highest accuracy of prediction when only one property is analyzed, is given by the isoelectric point, followed by mass and hydrophobicity.

When two properties are considered simultaneously, it was found that the combination of mass and the torsion angle is the most important in determining the presence of lysine glycation. As was outlined in previous research (Que-Salinas et al., 2022), this finding demonstrates that the occurrence of glycation results from the combination of several physical factors. This observation was already raised by Sjoblom et al. (2018) in suggesting some properties that could have considerable weight for arginine glycation, such as the clustering of acidic residues decreasing glycation levels while the presence of a single negative charge may be important for glycation success. However, it is important to note that, from the present investigation, and as noted in the three case studies, this does not imply that combining all the physical information available will result in an improved estimation of glycation likelihood.

Several approaches focused on identifying the presence or absence of glycation of proteinaceous lysine residues utilizing methodologies based on machine learning. The present work differs from these in a meaningful way, i.e., how the database is processed to apply machine learning techniques. In most of the previous studies, the database of thousands of amino acid sequences, once filtered, had been subjected to algorithms for the extraction of properties that allow the determination of the presence or absence of glycation, such as amino acid composition, encoding based on grouped weight or the K-nearest neighbor feature (e.g., Glypre (Zhao et al., 2017), PredGly (Yu et al., 2019), BERT-Kgly (Y. Liu et al., 2022)). Our work does not employ any of these algorithms but instead focuses on replacing the sequence of letters representing the amino acids by quantitative values of their different physical properties. Because of this, it is not possible to make a direct comparison of the results of this project with other studies regarding the presence of glycation in lysine for an independent test. A correct comparison (such as the one performed by Y. Liu et al. (2022)) implies that each of the models to be compared must be applied to the same database. Because of this difficulty, it should be added that not all the models developed for the present line of research are available or work well, as specified in several studies (Basith et al., 2021; Y. Liu et al., 2022; Yu et al., 2019). To summarize, an important consideration regarding the methodologies presented in the field is that artificial intelligence and, specifically, models based on artificial neural networks have gained significant strength and prominence in the last decade. In this regard, from the most current studies on the lysine glycation site, including the present work, recurrent neural networks based on LSTM algorithms represent one of the most promising methodologies.

Comparing the present results for lysine glycation with previous research (Que-Salinas et al., 2022) for arginine glycation, it is relevant to establish the cause of an apparent discrepancy in the results of both works. For lysine glycation the most determining physical property was IEP followed by Mass (Fig. 3 and 4), whereas in Que-Salinas et al. (2022) IEP was among the weakest performers. In contrast, the mass had an average performance. The following considerations explain this contrasting result: firstly, although the focus in both works is similar, which is to discern which physical properties have greater weight in the glycation process, the current work is focused on lysine, while in Que-Salinas et al. (2022), the study is focused on arginine. It is reasonable that different physical properties are the ones that influence with greater weight on the glycation process of different amino acids. While the previous work focuses on estimating the value of the probability of glycation from the database published by Sjoblom et al. in 2018, the present project focuses on classifying the presence or absence of glycation from the CPLM 4.0 database, so the approach of both methodologies is substantially different.

More studies are needed to contrast these results in determining the effect of the physical properties of amino near the glycation site on the glycation process. In summary, the properties of neighboring amino acids that play a significant role in the glycation process of lysine residues in proteins are both isoelectric point and mass.

## 5. Conclusions

This work outlines a comprehensive conceptual framework for predicting the susceptibility of proteinaceous lysine residues to glycation. A dataset of protein lysine modifications, filtered to exclude redundancy using the CD-HIT method, has been utilized. The database used is relatively large but does not contain quantitative data. It is also heterogeneous regarding methods that various laboratories have utilized to measure glycation in diverse biological systems. This contrasts with the relatively small quantitative data set on arginine glycation of short synthetic peptides reported by Sjoblom et al. (2018), which formed the basis of our previous article (Que-Salinas et al., 2022). Employing an RNN for lysine glycation classification allowed the identification of properties that are suggested to play an important role in lysine glycation. Specifically, by utilizing the IEP as the sole physical property for peptide characterization, a 59.6% accuracy in predicting lysine glycation was achieved. Furthermore, integrating two properties, Mass and ToA, increased the accuracy to 59.9%. Employing all eight properties led to a slightly reduced accuracy of 59.4%. The results obtained reflect the relevant relationships between the physical properties and the glycation process, meaning that the sequential structure of the peptide plays an important role. Our approach is designed to contribute to the existing landscape of the lysine-residue glycation estimation algorithms and to expand and enhance this landscape substantively. Although a perfect tool for unfailing predictions does not exist yet, our approach using the physical properties of amino acids neighboring the glycation site to determine its modification probability is pointing to a promising direction.

## References

Ahmed, N., Babaei-Jadidi, R., Howell, S. K., Beisswenger, P. J., & Thornalley, P. J. (2005). Degradation products of proteins damaged by glycation, oxidation and nitration in clinical type 1 diabetes. Diabetologia, 48(8), 1590–1603. 10.1007/s00125-005-1810-7

Basith, S., Hasan, M. M., Lee, G., Wei, L., & Manavalan, B. (2021). Integrative machine learning framework for the identification of cell-specific enhancers from the human genome. Briefings in Bioinformatics, 22(6), bbab252. 10.1093/bib/bbab252

Chandriah, K. K., & Naraganahalli, R. V. (2021). RNN / LSTM with modified Adam optimizer in deep learning approach for automobile spare parts demand forecasting. Multimedia Tools and Applications, 80(17), 26145–26159. 10.1007/s11042-021-10913-0

Ho, J. M. L., Miller, C. A., Smith, K. A., Mattia, J. R., & Bennett, M. R. (2021). Improved pyrrolysine biosynthesis through phage assisted non-continuous directed evolution of the complete pathway. Nature Communications, 12(1), 3914. 10.1038/s41467-021-24183-9

Huang, Y., Niu, B., Gao, Y., Fu, L., & Li, W. (2010). CD-HIT Suite: A web server for clustering and comparing biological sequences. *Bioinformatics (Oxford*, England), 26(5), 680–682. 10.1093/bioinformatics/btq003

Islam, M. M., Saha, S., Rahman, M. M., Shatabda, S., Farid, D. M., & Dehzangi, A. (2018). iProtGly-SS: Identifying protein glycation sites using sequence and structure based features. *Proteins: Structure*, Function, and Bioinformatics, 86(7), 777–789. 10.1002/prot.25511

Jia, J., Wu, G., & Li, M. (2024). iGly-IDN: Identifying Lysine Glycation Sites in Proteins Based on Improved DenseNet. Journal of Computational Biology: A Journal of Computational Molecular Cell Biology, 31(2), 161–174. 10.1089/cmb.2023.0112

Johansen, M. B., Kiemer, L., & Brunak, S. (2006). Analysis and prediction of mammalian protein glycation. Glycobiology, 16(9), 844–853. 10.1093/glycob/cwl009

Ju, Z., Sun, J., Li, Y., & Wang, L. (2017). Predicting lysine glycation sites using bi-profile bayes feature extraction. Computational Biology and Chemistry, 71, 98–103. 10.1016/j.compbiolchem.2017.10.004

Jurman, G., Riccadonna, S., & Furlanello, C. (2012). A Comparison of MCC and CEN Error Measures in Multi-Class Prediction. PLoS ONE, 7(8), e41882. 10.1371/journal.pone.0041882

Kumar Pasupulati, A., Chitra, P. S., & Reddy, G. B. (2016). Advanced glycation end products mediated cellular and molecular events in the pathology of diabetic nephropathy. Biomolecular Concepts, 7(5–6), 293–309. 10.1515/bmc-2016-0021

Liu, Y., Gu, W., Zhang, W., & Wang, J. (2015). Predict and Analyze Protein Glycation Sites with the mRMR and IFS Methods. BioMed Research International, 2015, 1–6. 10.1155/2015/561547

Liu, Y., Liu, Y., Wang, G.-A., Cheng, Y., Bi, S., & Zhu, X. (2022). BERT-Kgly: A Bidirectional Encoder Representations From Transformers (BERT)-Based Model for Predicting Lysine Glycation Site for *Homo sapiens*. Frontiers in Bioinformatics, 2:834153. 10.3389/fbinf.2022.834153

Liu, Z., Cao, J., Gao, X., Zhou, Y., Wen, L., Yang, X., Yao, X., Ren, J., & Xue, Y. (2011). CPLA 1.0: An integrated database of protein lysine acetylation. Nucleic Acids Research, 39(suppl_1), D1029–D1034. 10.1093/nar/gkq939

Liu, Z., Wang, Y., Gao, T., Pan, Z., Cheng, H., Yang, Q., Cheng, Z., Guo, A., Ren, J., & Xue, Y. (2014). CPLM: A database of protein lysine modifications. Nucleic Acids Research, 42(D1), D531– D536. 10.1093/nar/gkt1093

Mercado-Uribe, H., Andrade-Medina, M., Espinoza-Rodríguez, J. H., Carrillo-Tripp, M., & Scheckhuber, C. Q. (2020). Analyzing structural alterations of mitochondrial intermembrane space superoxide scavengers cytochrome-c and SOD1 after methylglyoxal treatment. PLOS ONE, 15(4), e0232408. 10.1371/journal.pone.0232408

Müller, M. M. (2018). Post-Translational Modifications of Protein Backbones: Unique Functions, Mechanisms, and Challenges. Biochemistry, 57(2), 177–185. 10.1021/acs.biochem.7b00861

Oya, T., Hattori, N., Mizuno, Y., Miyata, S., Maeda, S., Osawa, T., & Uchida, K. (1999). Methylglyoxal Modification of Protein. Journal of Biological Chemistry, 274(26), 18492– 18502. 10.1074/jbc.274.26.18492

Phillips, S. A., & Thornalley, P. J. (1993). The formation of methylglyoxal from triose phosphates: Investigation using a specific assay for methylglyoxal. European Journal of Biochemistry, 212(1), 101–105. 10.1111/j.1432-1033.1993.tb17638.x

Que-Salinas, U., Martinez-Peon, D., Reyes-Figueroa, A. D., Ibarra, I., & Scheckhuber, C. Q. (2022). On the Prediction of In Vitro Arginine Glycation of Short Peptides Using Artificial Neural Networks. Sensors, 22(14), 5237. 10.3390/s22145237

Rabbani, N., & Thornalley, P. J. (2014). Measurement of methylglyoxal by stable isotopic dilution analysis LC-MS/MS with corroborative prediction in physiological samples. Nature Protocols, 9(8), 1969–1979. 10.1038/nprot.2014.129

Rabbani, N., & Thornalley, P. J. (2015). Dicarbonyl stress in cell and tissue dysfunction contributing to ageing and disease. Biochemical and Biophysical Research Communications, 458(2), 221–226. 10.1016/j.bbrc.2015.01.140

Rabbani, N., & Thornalley, P. J. (2021). Protein glycation – biomarkers of metabolic dysfunction and early-stage decline in health in the era of precision medicine. Redox Biology, 42, 101920. 10.1016/j.redox.2021.101920

Rabuñal, J. R., & Dorado, J. (Eds.). (2006). Artificial Neural Networks in Real-Life Applications: IGI Global. 10.4018/978-1-59140-902-1

Reddy, H. M., Sharma, A., Dehzangi, A., Shigemizu, D., Chandra, A. A., & Tsunoda, T. (2019). GlyStruct: Glycation prediction using structural properties of amino acid residues. BMC Bioinformatics, 19(13), 547. 10.1186/s12859-018-2547-x

Schalkwijk, C. G., & Stehouwer, C. D. A. (2020). Methylglyoxal, a Highly Reactive Dicarbonyl Compound, in Diabetes, Its Vascular Complications, and Other Age-Related Diseases. Physiological Reviews, 100(1), 407–461. 10.1152/physrev.00001.2019

Scheckhuber, C. Q. (2019). Studying the mechanisms and targets of glycation and advanced glycation end-products in simple eukaryotic model systems. International Journal of Biological Macromolecules, 127, 85–94. 10.1016/j.ijbiomac.2019.01.032

Sjoblom, N. M., Kelsey, M. M. G., & Scheck, R. A. (2018). A Systematic Study of Selective Protein Glycation. Angewandte Chemie International Edition, 57(49), 16077–16082. 10.1002/anie.201810037

Williams, G. J., & Thorson, J. S. (2008). Natural Product Glycosyltransferases: Properties and Applications. In E. J. Toone (Ed.), Advances in Enzymology—And Related Areas of Molecular Biology (Vol. 76, pp. 55–119). John Wiley & Sons, Inc. 10.1002/9780470392881.ch2

Xu, H., Zhou, J., Lin, S., Deng, W., Zhang, Y., & Xue, Y. (2017). PLMD: An updated data resource of protein lysine modifications. Journal of Genetics and Genomics, 44(5), 243–250. 10.1016/j.jgg.2017.03.007

Xu, Y., Li, L., Ding, J., Wu, L.-Y., Mai, G., & Zhou, F. (2017). Gly-PseAAC: Identifying protein lysine glycation through sequences. Gene, 602, 1–7. 10.1016/j.gene.2016.11.021

Yu, J., Shi, S., Zhang, F., Chen, G., & Cao, M. (2019). PredGly: Predicting lysine glycation sites for *Homo sapiens* based on XGboost feature optimization. Bioinformatics, 35(16), 2749–2756. 10.1093/bioinformatics/bty1043

Zhang, W., Tan, X., Lin, S., Gou, Y., Han, C., Zhang, C., Ning, W., Wang, C., & Xue, Y. (2022). CPLM 4.0: An updated database with rich annotations for protein lysine modifications. Nucleic Acids Research, 50(D1), D451–D459. 10.1093/nar/gkab849

Zhang, Y., Sun, Z., Jia, J., Du, T., Zhang, N., Tang, Y., Fang, Y., & Fang, D. (2021). Overview of Histone Modification. In D. Fang & J. Han (Eds.), Histone Mutations and Cancer (Vol. 1283, pp. 1–16). Springer Singapore. 10.1007/978-981-15-8104-5_1

Zhao, X., Zhao, X., Bao, L., Zhang, Y., Dai, J., & Yin, M. (2017). Glypre: In Silico Prediction of Protein Glycation Sites by Fusing Multiple Features and Support Vector Machine. Molecules, 22(11), 1891. 10.3390/molecules22111891

