## Supplementary material for "Addressing the problem of lysine glycation prediction in proteins via Recurrent Neural Networks"

Ulices Que-Salinas[1], Dulce Martinez-Peon[2], Gerardo Maximiliano Mendez[2], P. Argüelles-Lucho[3], Angel D. Reyes-Figueroa[4,5], Christian Quintus Scheckhuber[6,*]

1 Centro de Ciencias de la Tierra, Universidad Veracruzana, Xalapa 91090, VER, México

2 Tecnológico Nacional de México/ Instituto Tecnológico de Nuevo León, Departamento de Ingeniería Eléctrica y Electrónica, Av. Eloy Cavazos 2001, Guadalupe, NL, México

3 Tecnológico Nacional de México, Instituto Tecnológico de Veracruz, Calzada Miguel Ángel de Quevedo 2779, Veracruz, Ver 91860, México

4 Consejo Nacional de Ciencia y Tecnología, Av. Insurgentes Sur 1582, Col. Crédito Constructor, Benito Juárez, México City 03940, DF, México

5 Centro de Investigación en Matemáticas Unidad Monterrey, Parque de Investigación e Innovación Tecnológica (PIIT), Av. Alianza Centro No. 502, Apodaca 66628, NL, México

6 Tecnológico de Monterrey, Escuela de Ingeniería y Ciencias, Av. Eugenio Garza Sada 2501 Sur, Col. Tecnológico, Monterrey 64700, NL, México

**Supplementary methods**

**Table S1. Numerical values corresponding to the eight physical properties of amino acids used in this study.** To form the vectors corresponding to each of the 6830 sequences of 31 amino acids, the letter which represents each amino acid is treated as a label that is exchanged for the numerical value based on the chosen physical property. The first column contains the names of the amino acids, while the next two tables contain their abbreviations. From the fourth column onwards, the numerical values of the 20 amino acids are presented for each of the eight physical properties.

| Full Name | Abbreviation | | Properties | | | | | | | |
|---|---|---|---|---|---|---|---|---|---|---|
| | 3 Letter | 1 Letter | Structure of the amino acid sequence (SoA) | Hydropathy (Hyd) | Mass (Mass) | Hydrophobicity (Hyp) | Polarizability (Pol) | Normalized van der Waals volume (vdW) | Torsion angle (ToA) | Isoelectric point (IEP) |
| Alanine | Ala | A | 1 | 1.8 | 89 | 0.31 | 0.05 | 1 | 4.76 | 6.11 |
| Arginine | Arg | R | -12 | -4.5 | 174 | -1.01 | 0.29 | 6.13 | 4.3 | 10.74 |
| Asparagine | Asn | N | -7 | -3.5 | 132 | -0.6 | 0.13 | 2.95 | 3.64 | 6.52 |
| Aspartate | Asp | D | -8 | -3.5 | 133 | -0.77 | 0.11 | 2.78 | 3.69 | 2.95 |
| Cysteine | Cys | C | 3 | 2.5 | 121 | 1.54 | 0.13 | 2.43 | 3.67 | 6.35 |
| Glutamine | Gln | Q | -9 | -3.5 | 146 | -0.22 | 0.18 | 3.95 | 4.54 | 5.65 |
| Glutamate | Glu | E | -10 | -3.5 | 147 | -0.64 | 0.15 | 3.78 | 5.48 | 3.09 |
| Glycine | Gly | G | 0 | -0.4 | 75 | 0 | 0 | 0 | 3.77 | 6.07 |
| Histidine | His | H | -6 | -3.2 | 155 | 0.13 | 0.23 | 4.66 | 2.84 | 7.69 |
| Isoleucine | Ile | I | 7 | 4.5 | 131 | 1.8 | 0.19 | 4 | 4.81 | 6.04 |
| Leucine | Leu | L | 5 | 3.8 | 131 | 1.7 | 0.19 | 4 | 4.79 | 6.04 |
| Lysine | Lys | K | -11 | -3.9 | 146 | -0.99 | 0.22 | 4.77 | 4.27 | 9.99 |
| Methionine | Met | M | 2 | 1.9 | 149 | 1.23 | 0.22 | 4.43 | 4.25 | 5.71 |
| Phenylalanine | Phe | F | 4 | 2.8 | 165 | 1.79 | 0.29 | 5.89 | 4.31 | 5.67 |
| Proline | Pro | P | -5 | -1.6 | 115 | 0.72 | 0.12 | 2.72 | 2,84 | 6.8 |
| Serine | Ser | S | -2 | -0.8 | 105 | -0.04 | 0.06 | 1.6 | 3.83 | 5.7 |
| Threonine | Thr | T | -1 | -0.7 | 119 | 0.26 | 0.11 | 2.6 | 3.87 | 5.6 |
| Tryptophan | Trp | W | -3 | -0.9 | 204 | 2.25 | 0.41 | 8.08 | 4.75 | 5.94 |
| Tyrosine | Tyr | Y | -4 | -1.3 | 181 | 0.96 | 0.3 | 6.47 | 4.3 | 5.66 |
| Valine | Val | V | 6 | 4.2 | 117 | 1.22 | 0.14 | 3 | 4.86 | 6.02 |

**Supplementary results**

The results are presented below in tables containing the quantitative values, as well as graphs in order to allow making comparisons. For case 1, each of the eight physical properties was selected separately to run the numerical algorithm, and the values corresponding to the five metrics were calculated from the results obtained by the RNN (Table S2). For case 2, two of the physical properties were selected to run the numerical algorithm and the values corresponding to the five metrics were calculated from the results obtained by the RNN (Table S3). For case 3, all the physical properties corresponding to case 1 were selected to run the numerical algorithm and the values corresponding to the five metrics were calculated from the results obtained by the RNN (Table S4).

**Table S2. The mean values of the classification results for one property (case 1).** The data have been sorted in descending order of accuracy. The values in bold text are the highest for each metric. Acc: accuracy, Pre: precision, Sen: sensitivity, Spe: specificity, MCC: Matthews Correlation Coefficient.

| Properties | Acc | Pre | Sen | Spe | MCC |
|---|---|---|---|---|---|
| **IEP** | **0.5961** | **0.5839** | 0.6399 | **0.5534** | 0.1947 |
| **Mass** | 0.5862 | 0.5572 | 0.8036 | 0.374 | **0.1963** |
| **Hyp** | 0.5841 | 0.5714 | 0.6391 | 0.5305 | 0.1713 |
| **vdW** | 0.5807 | 0.5579 | 0.7568 | 0.4087 | 0.1775 |
| **Hyd** | 0.5785 | 0.553 | 0.7695 | 0.392 | 0.175 |
| **Pol** | 0.5782 | 0.5551 | 0.7661 | 0.3947 | 0.1752 |
| **SoA** | 0.5761 | 0.5507 | 0.7715 | 0.3855 | 0.1716 |
| **ToA** | 0.5761 | 0.5486 | **0.8345** | 0.3237 | 0.1879 |

**Table S3. Performance of classification for the combination of two properties (case 2).** The data have been sorted in descending order of accuracy. The values in bold text are the highest for each metric. Acc: accuracy, Pre: precision, Sen: sensitivity, Spe: specificity, MCC: Matthews Correlation Coefficient.

| Two properties | Acc | Pre | Sen | Spe | MCC |
|---|---|---|---|---|---|
| Mass-ToA | **0.59935** | 0.582911 | 0.675202 | 0.525296 | 0.204667 |
| SoA-Hyp | 0.59785 | 0.577849 | 0.696761 | 0.501285 | 0.203254 |
| SoA-IEP | 0.59405 | 0.583299 | 0.630162 | 0.558794 | 0.190302 |
| Mass-Hyp | 0.59385 | 0.570103 | 0.74413 | 0.447134 | 0.203743 |
| Hyp-Pol | 0.59315 | 0.565774 | 0.782287 | 0.408498 | **0.209863** |
| Hyp-vdW | 0.58965 | 0.565676 | 0.771356 | 0.412253 | 0.202146 |
| ToA-IEP | 0.5896 | 0.574368 | 0.660425 | 0.520455 | 0.184202 |
| Hyp-IEP | 0.58835 | 0.574858 | 0.64251 | 0.535474 | 0.179306 |
| Mass-IEP | 0.58785 | **0.58331** | 0.582389 | 0.593182 | 0.175917 |
| Hyd-IEP | 0.58685 | 0.574435 | 0.633907 | 0.540909 | 0.176063 |
| Mass-vdW | 0.586 | 0.557809 | 0.787247 | 0.389526 | 0.192511 |
| Hyd-Hyp | 0.585 | 0.568719 | 0.664474 | 0.507411 | 0.174965 |
| vdW-ToA | 0.584 | 0.569207 | 0.676215 | 0.493972 | 0.175731 |
| Hyd-Pol | 0.58295 | 0.55725 | 0.771964 | 0.398419 | 0.185834 |
| Hyd-Mass | 0.5826 | 0.555974 | 0.784312 | 0.385672 | 0.187493 |
| Pol-IEP | 0.581 | 0.570902 | 0.616802 | 0.546047 | 0.164265 |
| SoA-Hyd | 0.57985 | 0.551462 | 0.802834 | 0.362154 | 0.184381 |
| SoA-vdW | 0.57965 | 0.551396 | 0.825405 | 0.339723 | 0.192076 |
| SoA-Pol | 0.57945 | 0.550346 | **0.826316** | 0.338439 | 0.191261 |
| SoA-Mass | 0.5792 | 0.552159 | 0.794028 | 0.369466 | 0.182328 |
| Hyd-vdW | 0.5785 | 0.550321 | 0.808198 | 0.354249 | 0.183056 |
| vdW-IEP | 0.57715 | 0.566692 | 0.615486 | 0.539723 | 0.156209 |
| Pol-ToA | 0.57685 | 0.562086 | 0.668117 | 0.487747 | 0.15987 |
| SoA-ToA | 0.5754 | 0.552194 | 0.75081 | **0.75081** | 0.168006 |
| Pol-vdW | 0.5722 | 0.550802 | 0.73836 | 0.40998 | 0.157936 |
| Mass-Pol | 0.57165 | 0.545927 | 0.802429 | 0.346344 | 0.167115 |

| | | | | | |
|---|---|---|---|---|---|
| Hyd-ToA | 0.5699 | 0.546433 | 0.766397 | 0.378063 | 0.159223 |
| Hyp-ToA | 0.56505 | 0.552098 | 0.641397 | 0.490514 | 0.134274 |

**Table S4. Performance of classification for the combination of all eight properties.** Acc: accuracy, Pre: precision, Sen: sensitivity, Spe: specificity, MCC: Matthews Correlation Coefficient.

| Properties | Acc | Pre | Sen | Spe | MCC |
|---|---|---|---|---|---|
| **All** | 0.5944 | 0.59 | 0.5925 | 0.5961 | 0.1903 |

For a detailed analysis of the results, the following figures S1 to S5 correspond to bar graphs showing the values obtained for each of the metrics calculated. For simplicity, the values of the same metric for cases 1 and 3 are shown in the same figure.

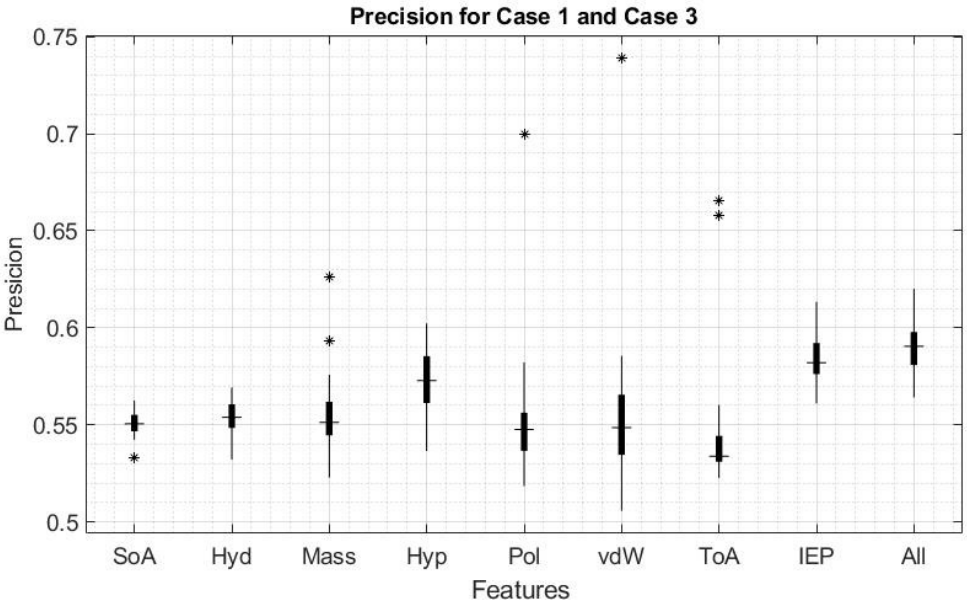


**Figure S1. Detailed analysis of the precision for case 1 and case 3 (All).** The x axis lists the eight analyzed properties. Median values are indicated by horizontal lines, the * symbol denotes outliers.

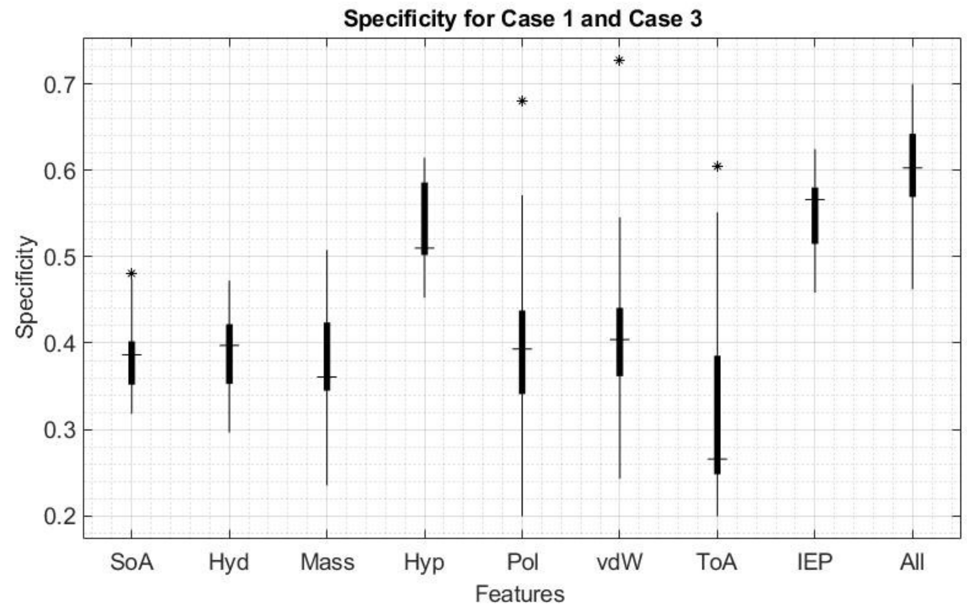


**Figure S2. Detailed analysis of the specificity for case 1 and case 3 (All).** The x axis lists the eight analyzed properties. Median values are indicated by horizontal lines, the * symbol denotes outliers.

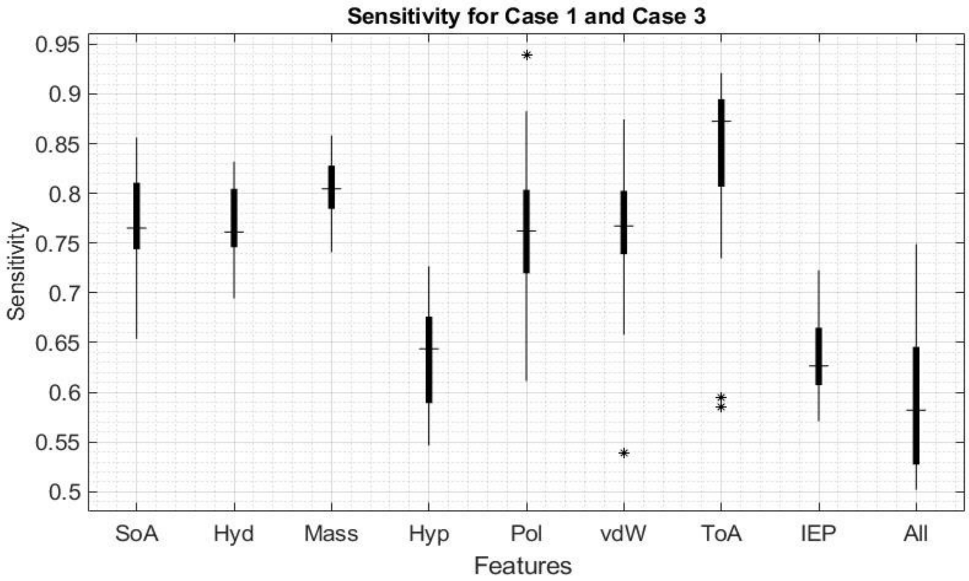


**Figure S3. Detailed analysis of the sensitivity for case 1 and case 3 (All).** The x axis lists the eight analyzed properties. Median values are indicated by horizontal lines, the * symbol denotes outliers.

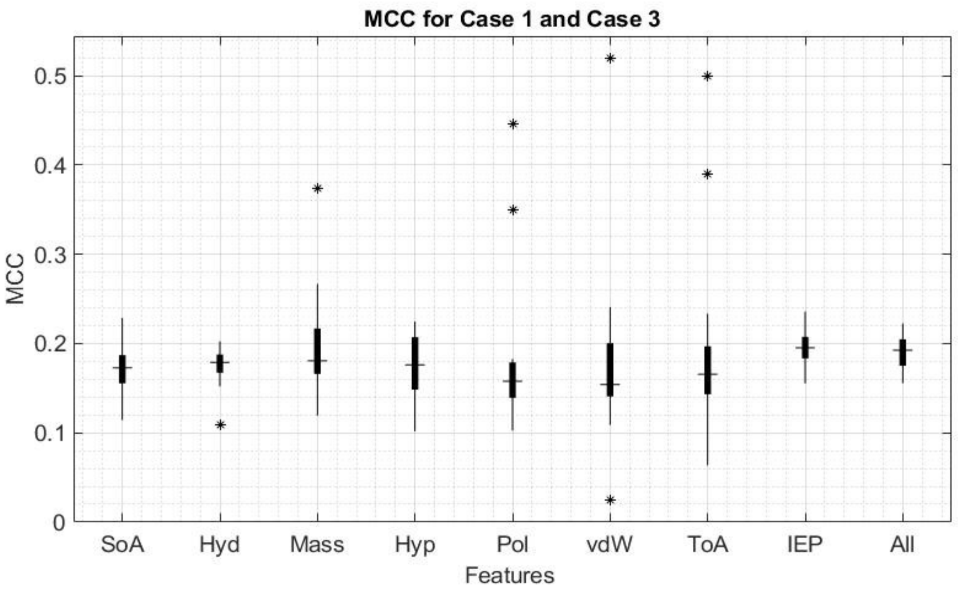


**Figure S4. Detailed analysis of the MCC for case 1 and case 3 (All).** The x axis lists the eight analyzed properties. Median values are indicated by horizontal lines, the * symbol denotes outliers.

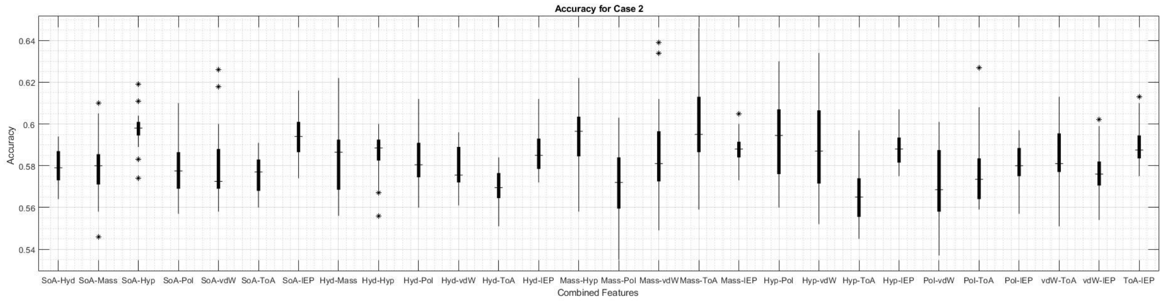


**Figure S5. Detailed analysis of the accuracy for case 2.** The x axis lists the combined properties. Median values are indicated by horizontal lines, the * symbol denotes outliers.
